# Structured behavioral data format: An NWB extension standard for task-based behavioral neuroscience experiments

**DOI:** 10.1101/2024.01.08.574597

**Authors:** Ryan Ly, Matthew Avaylon, Michael Wulf, Adam Kepecs, Oliver Rübel

## Abstract

Understanding brain function necessitates linking neural activity with corresponding behavior. Structured behavioral experiments are crucial for probing the neural computations and dynamics underlying behavior; however, adequately representing their complex data is a significant challenge. Currently, a comprehensive data standard that fully encapsulates task-based experiments, integrating neural activity with the richness of behavioral context, is lacking. We designed a data model, as an extension to the NWB neurophysiology data standard, to represent structured behavioral neuroscience experiments, spanning stimulus delivery, timestamped events and responses, and simultaneous neural recordings. This data format is validated through its application to a variety of experimental designs, showcasing its potential to advance integrative analyses of neural circuits and complex behaviors. This work introduces a comprehensive data standard designed to capture and store a spectrum of behavioral data, encapsulating the multifaceted nature of modern neuroscience experiments.

## 1. Introduction

The primary function of the brain is to generate behavior, making it essential to simultaneously study both neuronal activity and behavior to comprehend the functioning of neural circuits. Living organisms constantly encounter and respond to myriads of behavioral contingencies. To probe these, researchers design behavioral tasks to investigate specific processes ranging from sensory decisions to working memory, yet standardizing these tasks is fraught with challenges. A principal difficulty lies in the vast array of potential behavioral responses and their variation across numerous behavioral contingencies.

In the laboratory, a task controller (e.g., BAABL [16]) creates the defined experimental environment (e.g., to present stimuli) to subjects and reacts to responses from subjects based on the programmed behavioral contingencies (**Fig.1.1a**). Behavioral contingencies state the if-then conditions that set the occasion for the potential occurrence of certain behavior and its consequences (“If you move to the correct side, you will get a drop of water.”). The subject in turn receives certain stimuli and executes corresponding behavioral responses. Both, the experimenter and subject, hence, exhibit physical and algorithmic components that interact with each other as part of the experiment (see **Fig.1.1b**). By creating an appropriate environment (a), controlling the behavioral contingencies (c), and observing a subject’s actions and responses (b), the experimenter can then infer the brain’s algorithm (d) to produce the observed response.

**Figure 1.1.**
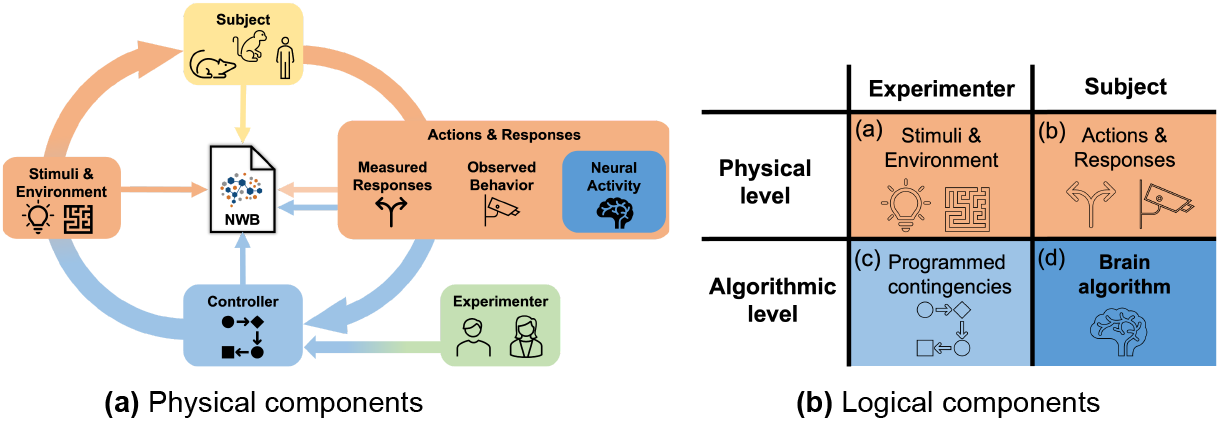
Components of a behavioral-task-based neurosciences experiment.

### Behavioral Control

There are limited commercial options available for controlling behavioral experiments. As a result, many labs rely on custom-made devices and software that do not adhere to a common, general program logic or data format. There are a few open-source hardware/software solutions for controlling behavioral experiment, e.g., Bpod [15], BAABL [16], ArControl [9], Bonsai [10], pyControl [11], HERBs [12], AutoPilot [14].

To computationally model behavioral tasks, many popular control software (e.g., BAABL, ArControl, and Bonsai) employ variants of the finite state machine (FSM) model to describe behavioral tasks. In this model, the program describes an abstract machine that can be in exactly one of a finite number of states at any given time. Transitions from one state to another are triggered based on some inputs, e.g., the subject pushing a button. An FSM is defined by: i) a finite list of states the system can be in, ii) the initial state the execution starts from, and iii) the inputs, outputs, actions, and logic that define the transitions between states. To better support the needs of behavioral experiments, behavioral control systems commonly expand (or relax) the FSM model, e.g., to support timers and parametrization of states. In the context of behavioral experiments, we then commonly distinguish further between inputs and transitions as the result of system logic (termed *Actions*) and inputs from the environment (termed *Events*), e.g., as a result of the subject selecting a choice port.

Even though different control systems vary in their implementations of this model, the FSM model provides us with a broadly applicable abstraction that allows us to effectively model the data generated by these systems. From a data-modeling perspective, the data generated by a behavioral task program can be described by: i) the *states* the system can be in, ii) the *actions* that can occur as part of system logic, and iii) the *events* that can be triggered by the subject (or environment). Note, it is beyond the scope of this manuscript to create a common language for defining behavioral tasks, but the focus of this paper is to define a standard for storing, sharing, and reusing data generated by behavioral experiments that use structured behavioral tasks.

### Data standards

In this work we build on and extend the Neurodata Without Borders (NWB) [1] data standard to support task-based behavioral neuroscience experiments. NWB is a community-driven data standard and software ecosystem for neurophysiology that provides neuroscientists with a common standard to share, archive, use, and build analysis tools for neurophysiology data. A growing community ecosystem of software tools for data management, analysis, visualization, and archiving are supporting NWB [2]. NWB supports formal extensions to the standard to enable users to integrate new data types or metadata and publish extensions for review [17, 18] via the NWB Extension Catalog [1,3].

The NWB data standard is designed to store a variety of neurophysiology data, including data from intracellular and extracellular electrophysiology experiments, optical physiology experiments, and tracking and stimulus data. For behavioral data, the core NWB standard supports storage of subject position data via *SpatialSeries*; *PupilTracking* and *EyeTracking* data; and behavioral events as generic *TimeSeries*. In addition, modern pose-estimation software, such as DeepLabCut [4] and SLEAP [5], can store the estimated positions of labeled body parts over time, the likelihood of each estimate, and metadata about the method in NWB files using the *ndx-pose* extension [6]. With this, NWB provides an ideal starting point for this work, because it already supports most data types required to capture the results from behavioral neuroscience experiments. However, critically, NWB currently does not support the description and capture of data and metadata about behavioral tasks and from the systems controlling the execution of behavioral tasks.

### Contributions

To rigorously describe and capture behavioral experiments and thereby enable progress towards understanding brain function, we need a data format that enables storage of all aspects of a behavioral experiment (**Fig 1.1b**) to (a) describe the environment and stimuli, (b) the subjects actions and responses, (c) the controller and observed parameters, as well as (d) the neural activity required for decoding the brain algorithm. The lack of a common standard to describe and store results from behavioral experiments impedes precise communication of behavioral task designs, sharing and re-analysis of data, and reproduction of experiments. To address this challenge we describe here with *ndx-structured-behavior* a formal extension to the NWB data standard to support the standardized description, storage, and sharing of measurements from behavioral experiments.

## 2. Method

*ndx-structured-behavior* [7] defines a formal extension to NWB for storing task programs and behavioral data. The extension supports explicit definition of: (i) key task metadata (e.g., available states, events, and actions) (**Sec. 2.1**), (ii) recorded times of the task execution (**Sec. 2.2**), and (iii) definition of the trial structure linked to the task execution details (**Sec. 2.3**). A key advantage of this approach is that it allows storage of both the behavioral data and acquired neural signals aligned in time together in a common data standard (**Sec. 2.4**). We have developed tools to convert data from the Bpod [7] and ArControl [8] data acquisition systems using our extension. While we here use BAABL as an example, the data model of the extension is not specific to BAABL nor Bpod.

### 2.1. Behavioral task description

The first main part of our *ndx-structured-behavior* extension focuses on describing metadata about structured tasks (**Fig. 2.1a**). All of the task metadata is organized in a *Task* module stored in the /*general* group as part of the NWB file hierarchy.

**Figure 2.1.**
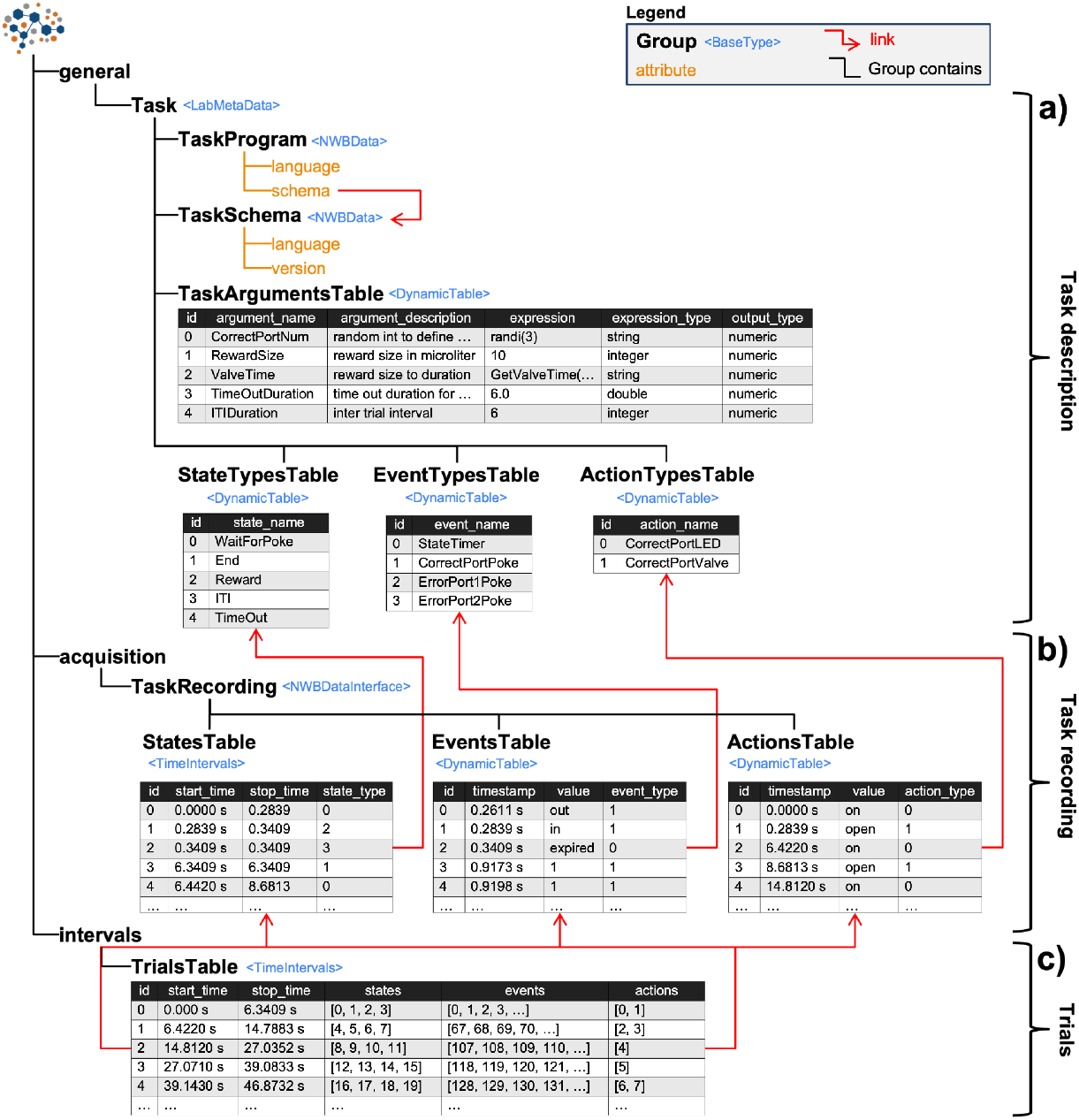
Overview of the schema of the extension.

Behavioral control systems use a broad range of methods for configuring tasks, from formal description languages (e.g., BAABL defined in XML), configuration files (e.g., in JSON), to custom code (e.g, Python scripts). To support this diversity of task program definitions, we support storage of arbitrary task descriptions as text dataset in the *TaskProgram* field. To facilitate interpretation, the *TaskProgram* is further described by the *language* (e.g. JSON, XML, Python etc.) used and an optional *TaskSchema* (e.g., a XML- or JSON-schema). This approach supports provenance and reuse, however, it does not allow users to easily interact and query key parameters of the task program due to the fact that the *TaskProgram* is usually specific to the control software and hardware.

To address this challenge, the *TaskArgumentsTable* is used to store the names and descriptions of all main parameters of the *TaskProgram* along with definitions of the *expression* used to define the parameters output values. Expressions may be simple constants (e.g., integers or floats) or complex functions, while the type of expression and type of parameter value generated are described by the *expression_type* and *output_type* columns, respectively.

In addition, we further abstract the definitions of states, events, and actions from the task program via the *StatesTypesTable, EventsTypesTable*, and *ActionsTypesTable*. The *TaskArgumentsTable, StatesTypesTable, EventsTypesTable*, and *ActionsTypesTable* are all defined using the NWB *DynamicTable* type, enabling users and tools to include additional metadata specific to the experiment or control system by adding custom columns to the tables. In practice, the metadata stored in these tables can be automatically extracted from parsing the *TaskProgram*, avoiding the need for manual data entry by the users.

### 2.2. Behavioral task recordings

Data acquired from the control system that describes the execution of the task program are then organized in the *TaskRecording* module and stored in the /*acquisition* group as part of the NWB file hierarchy.

In accordance with the permissible values in their respective types tables, the *StatesTable, EventsTable*, and *ActionsTable* store the timings of states, events, and actions during the experiment when running the *TaskProgram*. Each row in these tables represents a single instance of a state, event, or action with rows being sorted by time. The *StatesTable* inherits from NWB’s *TimeIntervals* type to effectively store start- and end-times for states in the experiment. The *EventsTable* and *ActionsTable* inherit from NWB *DynamicTable* to store timestamps of events and actions, as these are commonly treated as instantaneous. To also support non-instantaneous, finite-time actions and events, the *EventsTable* and *ActionsTable* define an optional *duration* column.

While executing a task program, the same state, event or action usually occurs many times. To define the type of state, event, and action, the tables include a *DynamicTableRegion* column to reference the corresponding row in *StatesTypesTable, EventsTypesTable*, and *ActionsTypesTable*, respectively. These references are automatically resolved for the user through PyNWB when converting to Pandas, ensuring quick data interpretability.

### 2.3. Trial structure

NWB supports annotation of the temporal structure of experiments via the *TimeIntervals* table type. *TimeIntervals* tables are stored in the */intervals* group in NWB, and are used to annotate: i) *epochs* describing experimental stages or sub-divisions of a single recording session, ii) *invalid_times* to demarcate time intervals that should be removed from analysis, and iii) *trials* describing repeated experimental events. In the case of trials, each row in the table represents a trial and stores the *start_time* and *stop_time* of the trial as a floating-point offset in seconds to the reference time of the NWB file. In addition, the *timeseries* column enables users to reference corresponding ranges of relevant *TimeSeries* recordings of, e.g. neural activity, stimuli, or behavior. Users may also include additional metadata about trials via custom columns.

Building on this structure, we are enhancing the trials table by defining columns to reference the corresponding behavioral task *states, events*, and *actions* that occurred during the trial (**Fig. 2.1c**). The *states, events*, and *actions* columns are defined using the *DynamicTableRegion* type in NWB, which allows us to reference rows in the *StatesTable, EventsTable*, and *ActionsTable*, respectively. Since in each trial, a variable number of states, events, and actions may occur, the columns are indexed in NWB. Via the PyNWB API, users can conveniently interact with this data structure as it automatically resolves the references to the *StatesTable, EventsTable*, and *ActionsTable*, enabling the users to easily access the relevant data about states, events, and actions related to a trial without having to manually navigate the different levels of the data structure in NWB.

### 2.4. Integration with neural data, stimuli, and free behavior

Our extension in combination with the existing capabilities of the NWB data standard, enables researchers for the first time to relate the: i) behavioral task descriptions and events, actions, and states data (via our *ndx-structured-behavior* extension), ii) pose estimates, e.g., from annotated videos and automatically tracked body parts (via ndx-pose [6]), iii) spatial navigation (NWB core), and iv) neural data (NWB core) with each other aligned in time via a single, coherent data standard. All timestamps in an NWB file are stored as floating-point offsets in seconds relative to a common reference time, making it easy to directly relate signals stored in an NWB file with each other, as well as align signals in time from different NWB files based on a single time offset based on the difference of the reference time used in the files.

## 3. Results

As a formal extension for NWB, users can use the standard PyNWB (Python) and MatNWB (Matlab) APIs for NWB to input, write, and read their data using *ndx-structured-behavior*. To demonstrate the application of our extension in practice, we have developed converters for storing recordings from BAABL (Behavioral tAsk Analysis & Building Language) using the Bpod controller and ArControl [8, 9] using our extension. Below we describe example uses of our extension to store the results from example behavioral task experiments.

### 3.1. Light chasing task

To illustrate the extension in practice we use a basic light chasing task implemented using BAABL (**Fig. 3.1**). The subject is in a behavior box with 3 ports (**Fig. 3.1a**). Each port consists of an IR-based poke detection with a white LED and valve to deliver a water reward via a spout. In each trial one port is randomly chosen as the correct port, while the other two ports are error ports. The LED is then activated in the correct port and the system is waiting for the subject to poke into a port. If the subject chooses the correct port with the LED, a predefined water reward is given. Afterwards, the system waits for a pre-defined, inter-trial-interval (ITI) duration before starting the next trial.

**Figure 3.1.**
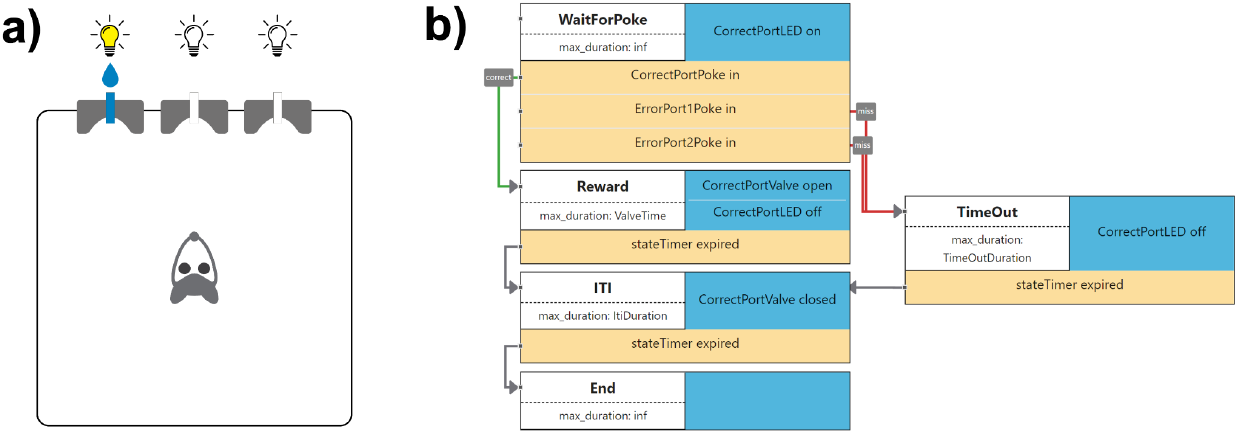
Overview of the light chasing task. **a)** Illustration of the subject in the behavior box with 3 ports. **b)** Illustration of the state-machine diagram for the task.

**Figure 3.2.**
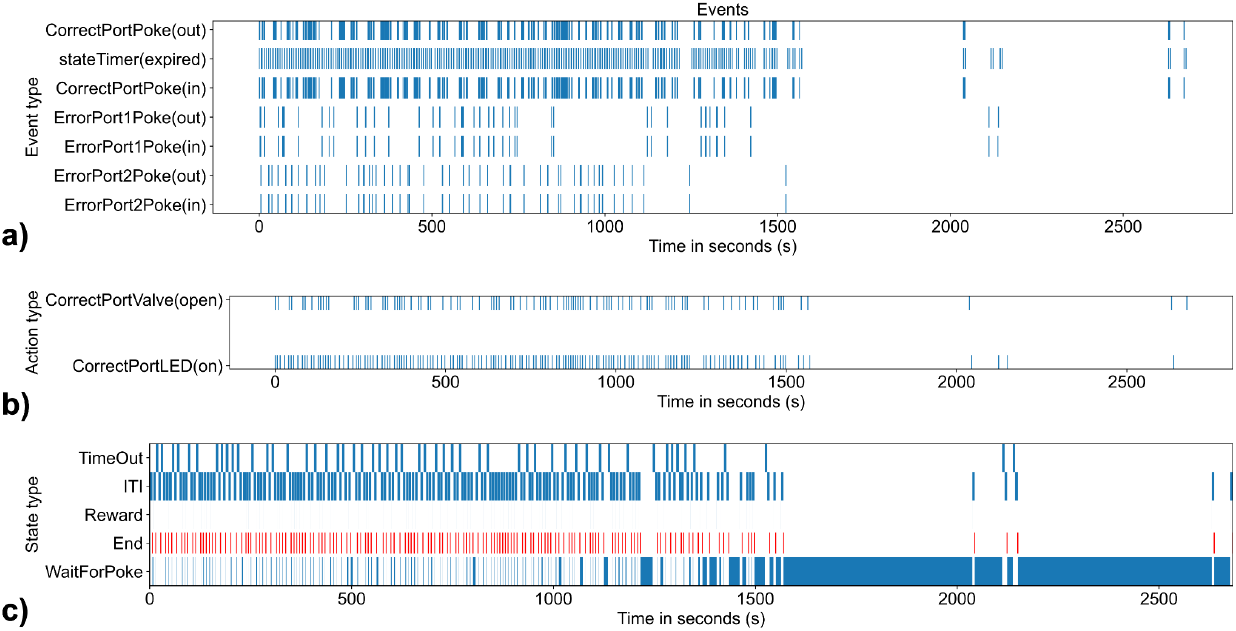
Time raster plot showing the **a)** events, **b)** actions, and **c)** states recorded from the BAABL control system.

**Figure 3.3.**
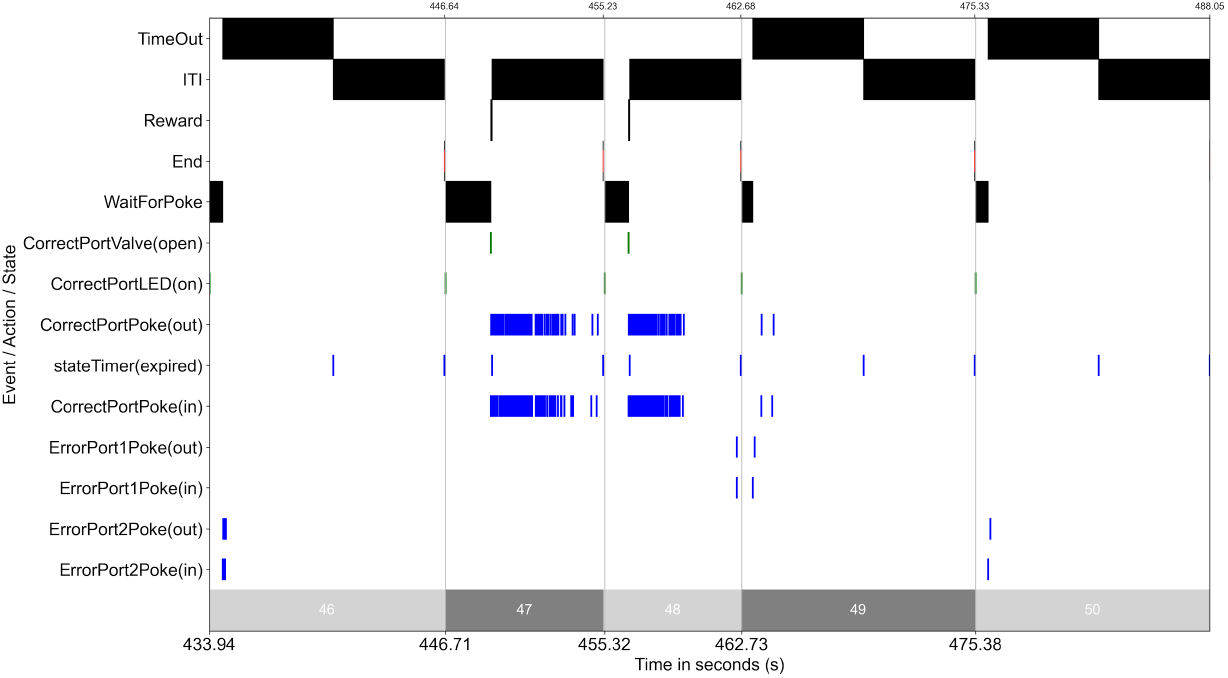
Time raster plot of the *TrialsTable*, showing trials 46 to 50 of the light chasing task behavior experiment, including all states (top), actions (center), and events (bottom) recorded from the BAABL control system.

**Figure 3.4.**
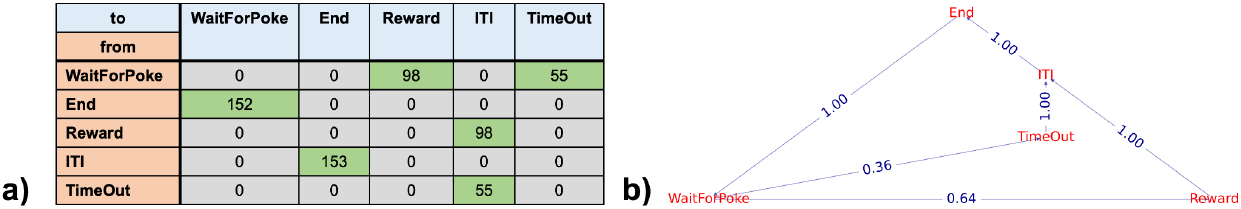
State transition analysis for the light chasing task example. **a)** Matrix plot showing the number of transitions from one state to another state during the experiment. **b)** Visualization of the transition matrix shown in (a) as a transition probability graph.

**Fig. 2.1** illustrates the layout of the light chasing task data in NWB recorded via BAABL using the BPod system. Metadata describing the task is stored in the **/general** group in NWB and is typically visualized using standard table displays, e.g., via Pandas.

To simplify inspection of events, actions, and states recorded from the control system, our extension provides convenient methods for generating time raster plots (**Fig. 3.2**). Acquired, event-specific information (e.g., timing and value) are retrieved from the *EventsTable* stored in the */acquisition* group, while metadata about the corresponding event type used for labeling are retrieved from the *EventTypesTable* in the */general* group (**Fig. 2.1**). Correspondingly the *ActionsTable* plus *ActionTypesTable* and *StatesTable* plus *StateTypesTable* are used to plot the time raster for actions (**Fig. 3.2b**) and states (**Fig. 3.2c**), respectively.

All information is ultimately combined in the *TrialsTable*, which describes the trial structure of the data. Using the *TrialsTable* makes it easy to identify and access all events, actions, and states recorded during a trial for further analysis. To support visual inspection of the *TrialsTable*, our extension supports plotting of trials as time rasters, showing all events, actions, and states during the select trials in a single view (**Fig. 3.3**).

While the task program defines the structure of possible state transitions, the actual state transitions recorded during the experiment are dependent directly on the subject’s behavior and choices. From the *StatesTable* (and *StatesTypesTable*) we can easily count the number of times the system transitions from one state to another (see **Fig.3.4a**). We can then normalize the counts to compute the probabilities of transitions between states, and visualize the result as a state-transition graph (see **Fig. 3.4b**). Here we see, e.g., that in 64% of all trials the subject received a reward and 36% of trials timed out. The lower success-rate here is due to the fact that this data was recorded during an earlier training stage of the subject, while the mouse was still learning the task and still acclimating to the behavioral setup (especially the reward ports).

## 4. Discussion

The *ndx-structured-behavior* data standard lays the foundation towards a comprehensive standard for describing the data from end-to-end behavioral neuroscience experiments. Integrated with NWB, *ndx-structured-behavior* enables researchers for the first time to store and relate the data from structured behavioral task recordings, pose estimates, spatial navigation and neural data with each other aligned in time via a single, coherent data standard. Since our approach builds on the established NWB data standard, users are able to use our extensions to publish their data via the DANDI archive [1, 14], providing an easy-to-use mechanism for publication and collaboration.

However, several key challenges remain to enable FAIR reuse of structured behavioral tasks and data that are beyond the scope of this effort. Task programs are currently being implemented via a wide range of methods and formats that are usually specific to the control software and hardware. This approach hinders reuse and reproduction of behavioral tasks and experiments, requiring access to the same control software and hardware or reimplementation of task programs, which is error-prone and hard due to the varying capabilities and designs of different control systems. BAABL describes a first major step in this direction by defining a hardware-agnostic, portable language for describing behavioral tasks. However, more effort is needed to harmonize and enable interoperability of task programs across control software/hardware solutions. Other key challenges are driven by gaps in the software ecosystem for structured behavioral tasks. For example, the community lacks tools to formally evaluate behavioral tasks prior to running experiments in the field. This leads to late detection of errors, hinders computational modeling of experiments, and promises to be an invaluable tool for computational evaluation of brain algorithm models derived from experiments.

## Acknowledgements

Research reported in this publication was supported by the National Institute of Mental Health under Award Number 7RF1MH120034 (PI: A. Kepecs) and National Institute of Neurological Disorders and Stroke under Award Number 5U24NS120057 (PI: O. Ruebel). Additional support for NWB was also provided by the Kavli Foundation.

We thank the diverse participants of the NWB hackathons and the global NWB user and developer community for their feedback and contributions. We thank the current and former members of the NWB Executive Board and NWB Technical Advisory Board for their guidance and support: Kristofer Bouchard, Bing Brunton, Elizabeth Buffalo, Anne Churchland, Loren Frank, Satrajit Ghosh, Adam Kepecs, Lydia Ng, Huib Mansvelder, Ueli Rutishauser, Karel Svoboda, Christof Koch, Friedrich Sommer, Markus Meister, Katrin Amunts, Saskia de Vries, Anna (Szonja) Weigl, Alessio Buccino, Yaroslav O. Halchenko, and Lawrence Niu.

## Disclaimer

This document was prepared as an account of work sponsored by the United States Government. While this document is believed to contain correct information, neither the United States Government nor any agency thereof, nor the Regents of the University of California, nor any of their employees, makes any warranty, express or implied, or assumes any legal responsibility for the accuracy, completeness, or usefulness of any information, apparatus, product, or process disclosed, or represents that its use would not infringe privately owned rights. Reference herein to any specific commercial product, process, or service by its trade name, trademark, manufacturer, or otherwise, does not necessarily constitute or imply its endorsement, recommendation, or favoring by the United States Government or any agency thereof, or the Regents of the University of California. The views and opinions of authors expressed herein do not necessarily state or reflect those of the United States Government or any agency thereof or the Regents of the University of California

## Availability of Code

The source for *ndx-structured-behavior* is available online at https://github.com/rly/ndx-structured-behavior. The source and data used in the example shown in Section 3.1 and the plots shown in Figure 3.2, 3.3, and 3.4 are included in the tutorial section of the repository at https://github.com/rly/ndx-structured-behavior/tree/main/docs/tutorials.

